# Accuracy of direct genomic values and methylation profile through genotype-by-LowPass sequencing using Nanopore technology

**DOI:** 10.1101/2023.01.15.523960

**Authors:** Oscar González-Recio, Adrián López-Catalina, Ramón Peiró-Pastor, Alicia Nieto-Valle, Monica Castro, Almudena Fernández

**Affiliations:** Dpt. Mejora Genética Animal. INIA-CSIC. Ctra La Coruña km 7.5, 28040 Madrid (Spain)

**Keywords:** Low Pass sequencing, low sequencing imputation, Genomic Selection, Polygenic Risk Score, Genomic values

## Abstract

Genotype-by-sequencing has been proposed as an alternative to SNP genotyping arrays in genomic selection to obtain a high density of markers along the genome. It requires a low sequencing depth to be cost effective, which may increase the error at the genotype assigment. Third generation Nanopore sequencing technology offers low cost sequencing and the possibility to detect genome methylation, which provides added value to genotype-by-sequencing. The aim of this study was to evaluate the performance of genotype-by-LowPass Nanopore sequencing for estimating the direct genomic value in dairy cattle, and the possibility to obtain methylation marks simultaneously. Latest Nanopore chemistry (LSK14 and Q20) achieved a modal base calling accuracy of 99.55 %, whereas previous kit (LSK109) achieved slightly lower accuracy (99.1 %). The direct genomic value accuracy from genotype-by-Low Pass sequencing ranged between 0.79 and 0.99, depending on the trait, with a sequencing depth as low as 2x and using the latest chemistry (LSK114). Lower sequencing depth led to biased estimates, yet with high rank correlations. The LSK109 and Q20 achieved lower accuracies (0.57-0.93). More than one million high reliable methylated sites were obtained, even at low sequencing depth, located mainly in distal intergenic (87 %) and promoter (5 %) regions. This study showed that the latest Nanopore technology can be use in a LowPass sequencing framework to estimate direct genomic values with high reliability. It may provided advantages in populations with no available SNP chip, or when a large density of markers with a wide range of allele frequencies is needed. In addition, Low Pass sequencing provided with nucleotide methylation status of >1 million nucleotides at ≥ 10x, which is an added value for epigenetic studies.

## Introduction

Advances in genotyping platforms over the past two decades have enabled the prediction of genetic value in individuals for the implementation of genomic selection in animal and plant populations. They also allowed the prediction of polygenic risk scores in human populations that predict the probability of suffering certain diseases. Initially, genotyping arrays consisted of a few hundreds or thousands SNPs, but improvements in the technology soon after allowed for the incorporation of hundreds of thousands of SNPs in genotyping arrays. Methods for genotype imputation have also contributed to the use of different genotyping platforms or different densities of genotyping arrays. A major disadvantage of SNP arrays is that their design is often based on few animals/populations, which limits their use in other populations not considered in the design. In addition, low frequency and rare variants are seldom included in the genotyping arrays, which may miss linkage disequilibrium with relevant causal variants of certains diseases and traits. More recently, genotype-by-sequencing has allowed capturing millions of variants along the genome. Genotype-by-sequencing techniques can be used to align DNA reads against a reference genome and detect polymorphic positions with bioinformatics tools throughout the genome, regardless of whether they have been previously detected or included in an array design. The precision of this genotype-by-sequencing is mainly determined by the sequencing depth. However, the limitations in precision at low sequencing depth can be compensated for by imputation strategies, as its more affordable cost allows sequencing many more individuals in the population, improving the statistical power of genomic selection and genomic studies (1). Detecting a larger number of variants at different minor allele frequencies helps to discover association signals in genome-wide association studies and estimate the genetic value of individuals with a similar precision as SNP chips 2, 3. Genetic imputation has also been applied to genotype-by-sequencing data, which needs to deal with artifact errors due to low depth or LowPass Sequencing (LPS). Some methods have already been proposed to palliate this limitation (1, 4–6).

Third-generation sequencing techniques such as Nanopore have been explored as an option for genomic selection using information at low sequencing depth (7). This technique allows for fast and low-cost sequencing at the expense of a higher error rate compared to Sanger or sequencing by synthesis. However, the latest Nanopore chemistry offers higher accuracy which may increase the accuracy of the prediction of genetic values using this technique. Additionally, Nanopore sequencing can simultaneously detect epigenetic modifications at the nucleotide level, and it is obtained at no additional cost. This information can be used in breeding programs and epigenetic studies in livestock and plants (8). Nanopore sequencing has already been used for pathogen identification, metagenomic studies, and the assembly of reference genomes. However, its higher sequencing error rate has discouraged its use for predicting the genetic value of individuals. Since the accuracy and yield of the technique has improved in recent years, along with its low cost, better portability, the ability to obtain modified bases, and specific bioinformatics tools, it is now more attractive for exploring its use in genomic prediction in a genotype-by-LPS framework. It is also an alternative tool to genomic research involving epigenetics.

## Materials and Methods

### A. Samples and DNA extraction

Blood samples were obtained from 32 Holstein calves during routinary practices in a commercial farm in the Northeast regions of Spain. These samples were obtained by a veterinarian during the routinary process for genomic evaluations within the official Holstein breeding program in Spain (https://www.conafe.com). One sample from each animal was sent to the official genotyping lab, and was genotyped using the Illumina EUROG MD genotyping microarray that contains approximately 62,000 markers. Another sample was sent to the departmente of animal breeding at INIA-CSIC, where DNA was extracted using the Monarch HMW DNA Extraction Kit for Cells Blood. This DNA was then prepared for sequencing.

### B. Sequencing

The purified DNA was sequenced in either a MinION or GridION device from Oxford Nanopore Technologies. The individual DNA libraries were prepared starting with 3 µg of DNA. Twelve samples were sequenced using the kit SQK-LSK109 (LSK109) in R9.4 flow cells, multiplexing 6 samples per flow cell. Other twelve samples were sequenced using the kit SQK-LSK110 (Q20) in R9.4 flow cells, also multiplexing 6 samples per flow cell. Finally, the remaining six samples were sequenced following the protocol from the kit SQK-LSK114 (LSK114) in R10.4.1 flow cells, multiplexing 2 samples per flow cell. Two samples from LSK109 were discarded for not yielding enough reads to start the bioinformatic analyses.

### C. Bioinformatic pipeline

Basecalling was performed with guppy toolkit version 6.4.2 using SUP mode. Reads with length <150bp or ONT quality score <10 were discarded. Remaining reads were aligned against Bos taurus reference genome (ARS-UCD1.2) using minimap2 aligner, with option `-ax map-ont` (9), a general-purpose alignment program to map DNA or long mRNA sequences. Coverage statistics were calculated with `samtools coverage` (10). After the alignment, the content and percentage of mismatches by read were computed. The CIGAR string (10) and the edit distance from the reference or number of mismatches per pair (nM tag value) from the alignment were used to extract the total length of insertions and deletions and single nucleotides mismatches for each read. The nM tag value is the sum of total mismatch score (TMS) and length of insertions and deletions. Thus, TMS was computed as:

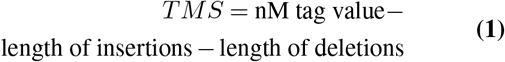

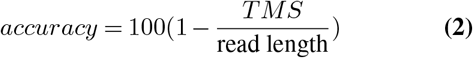

The aim of this study was to determine the accuracy of low-pass sequencing (LPS) using Nanopore technology in terms of basecalling, imputation, and prediction of genetic merit in comparison to SNP genotyping arrays within a genomic selection framework. Both older and more recent Nanopore chemistries were compared, and the potential to include epigenetic information was also evaluated.

Then, variants were called using `Clair3 v0.1-r11` (11). Variants with sequencing depth *≤*2 were discarded for downstream analysis unless the variant was equal to the alternate allele in the reference genome. A heterozygous position was called if the allele frequency was larger than 0 and lower than 90 %. The resulting variants were then imputed to whole genome sequencing using the 1000 bull genomes project (12) and `Beagle` version 5.2 (13). We kept those common variants in the Illumina Bovine50K beadchip that were included in the official genomic evaluations of Milk Yield (MY), Fat Yield (FY) and Protein Yield (PY) frome the Spanish Holstein Association (CONAFE).

### D. Computing direct genomic values

Direct genomic values (DGV_*it*_) for each individual *i* and trait (*t*=MY,FY,PY) (either from SNP beadchips or LPS) were calculated as:

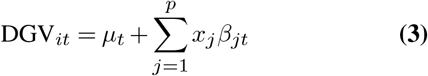

where *µ*_*t*_ is some intercept value specific for each trait, *x*_*j*_ is either the SNP genotype or the dosage allele (DA) from imputed ONT sequencing, and *β*_*jt*_ is the allele substitution effect for SNP *j* and trait *t* provided by CONAFE.

### E. Detection of modified based

Modified bases (5mc) were extracted from samples sequenced with LSK114 kit. Methylation marks were detected from bam files using `modbam2bed` tool provided by Nanopore software (14). Genetic features and coordinates were annotated using the R package `ChIPseeker` (15). Promoter regions were called using the function `getPromoter` using the transcription annotated genome for Bos Taurus and the annotation package `org.Bt.eg.db`. The transcription start site (TSS) region was defined as -3000 to 3000 base pairs from the transcription start site. Sequencing depth thresholds of 4x, 7x and 10x were compared to determine the variation in the genetic features lost when establishing a more stringent filter.

The genetic feature in which the methylation marks are located were called using the `plotAnnoBar` function. Then, heatmaps depicting the distribution of methylation marks in the promoter regions were obtained using the tagHeatmap function.

## Results

### F. Descriptive summary

A summary of the samples kept after quality control is shown in Table 1. The kit LSK109 showed higher yield than Q20, which translated into a higher average sequencing depth (0.6x vs 0.4x) and a larger number of called variants (221k vs 142k). Samples sequenced with the LSK114 kit showed a higher average sequencing depth (2.1x *±* 0.4x SD) and a larger initial number of variants (1635k *±* 455k SD). Improved yield from LSK114 was partially determined because only two samples were multiplexed per flowcell. However, it is equivalent to a 0.8x sequencing depth if six samples per flow cell would have been multiplexed as in LSK109 and Q20 kits. This circumstance is evaluated below to evaluate LSK114 under lower sequencing depth.

**Table 1.** Sequencing depth and number of variants left to start imputation after filtering for each sample

|  | **kit** | **Sequencing Depth** | **Number of variants** |
| --- | --- | --- | --- |
| Sample 1 | LSK109 | 0.36 | 69,700 |
| Sample 2 | LSK109 | 0.51 | 139,412 |
| Sample 3 | LSK109 | 0.41 | 84,872 |
| Sample 5 | LSK109 | 0.54 | 143,414 |
| Sample 6 | LSK109 | 0.54 | 151,828 |
| Sample 8 | LSK109 | 0.61 | 169,916 |
| Sample 9 | LSK109 | 0.45 | 108,916 |
| Sample 10 | LSK109 | 1.04 | 495,019 |
| Sample 11 | LSK109 | 0.88 | 361,627 |
| Sample 12 | LSK109 | 1.07 | 490,223 |
| Sample 13 | Q20 | 0.41 | 115,022 |
| Sample 14 | Q20 | 0.30 | 65,840 |
| Sample 15 | Q20 | 0.38 | 93,316 |
| Sample 16 | Q20 | 0.31 | 67,232 |
| Sample 17 | Q20 | 0.33 | 76,340 |
| Sample 18 | Q20 | 0.36 | 91,323 |
| Sample 19 | Q20 | 0.86 | 419,787 |
| Sample 20 | Q20 | 0.47 | 145,412 |
| Sample 21 | Q20 | 0.74 | 321,066 |
| Sample 22 | Q20 | 0.31 | 74,607 |
| Sample 23 | Q20 | 0.49 | 140,729 |
| Sample 24 | Q20 | 0.38 | 94,729 |
| Sample 25 | LSK114 | 2.43 | 1,924,878 |
| Sample 26 | LSK114 | 2.1 | 1,533,807 |
| Sample 27 | LSK114 | 1.87 | 1,311,229 |
| Sample 28 | LSK114 | 1.92 | 1,375,101 |
| Sample 29 | LSK114 | 1.78 | 1,246,664 |
| Sample 30 | LSK114 | 2.93 | 2,420,455 |

### G. Variant calling accuracy

Basecalling accuracy from each sequencing kit is depicted in Figure 1. Median accuracy was 98.5%, 98.7%, and 99.0% for the LSK109, Q20, and LSK114 kits, respectively. Mode accuracy was 99.1%, 99.6%, and 99.5% for the LSK109, Q20, and LSK114 kits, respectively. It must be noted that this is a down-limit accuracy because it was calculated against the reference genome, so true variants are incorrectly counted as errors. Nonetheless, a significant number of reads showed basecalling accuracies below 95%. These reads may introduce error variants in downstream analysis.

**Fig. 1.**
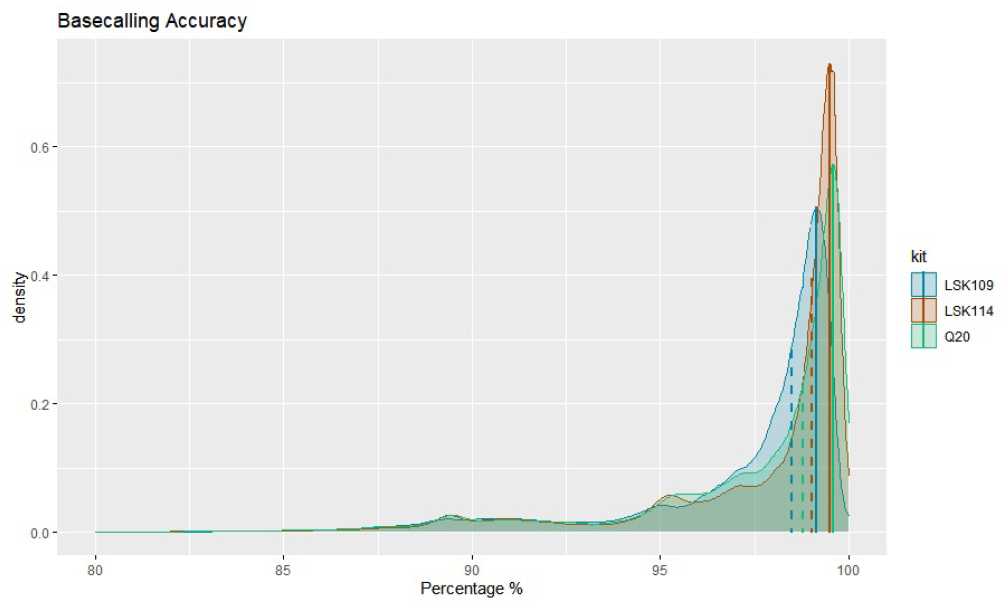
Density plot of the basecalling accuracy for each sequencing analysed, measured as equation (2). Mode value from each kit is depicted as a vertical solid line. Median value from each kit is depicted as a dashed vertical line

### H. Imputation accuracy

After imputing called variants to whole genome sequencing, the imputed variants were compared to the genotypes from the SNP array. Figure 2 shows a high degree of concordance between the imputed variant from LPS and the genotype. Lower agreement was observed for heterozygous genotypes when the LSK109 and Q20 kits were used. In these cases, imputation was less accurate. Samples sequenced with the LSK114 kit were accurately imputed, although wider ranges of DA were observed for homozygous SNPs when LPS variants were imputed from sequence depths as low as 0.5x. In contrats to older chemistry, more accurate DA was imputed from LSK114 even for heterozygote genotypes and at similar sequencing depths similar 0.5x.

**Fig. 2.**
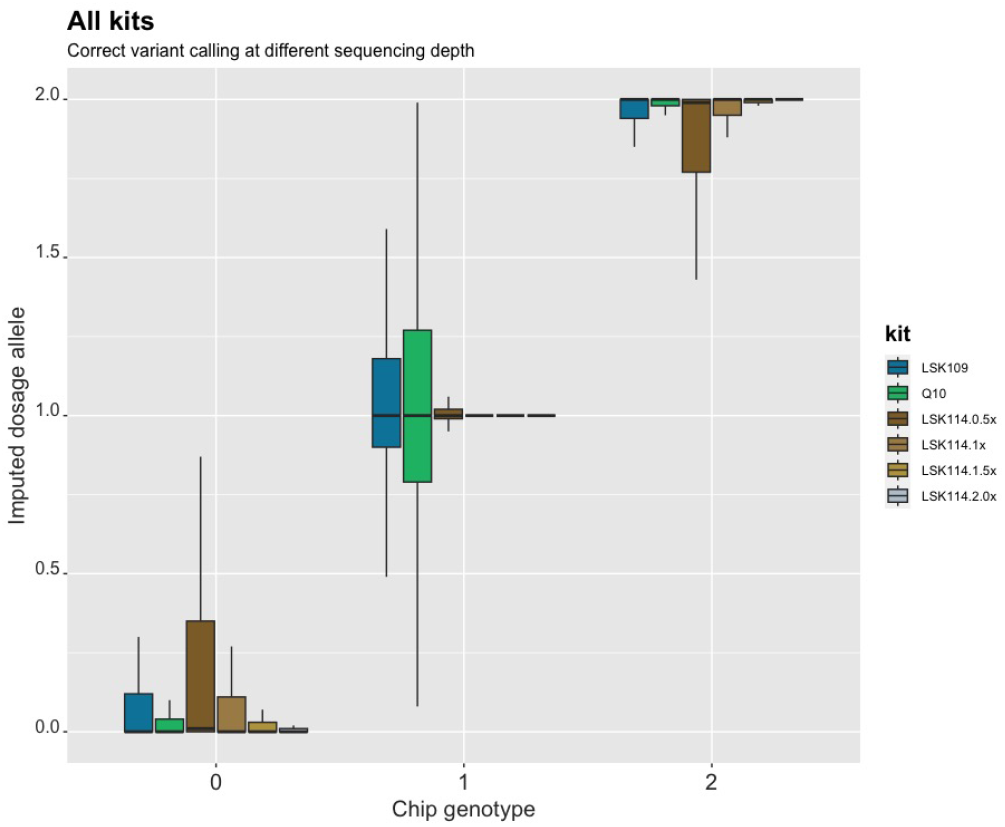
Imputed dosage allele obtained after LPS according to the SNP genotype code. Each kit and sequencing depth is depicted in different color.

Commonly, heterozygous genotypes are called for 0.8<DA<1.2. The percentage of correct and miscalled genotypes from LSK114 is shown in Figure 3 at different sequencing depths. A larger amount of corrected calls were imputed for homozygous positions ranging from 85.2% at a sequencing depth of 0.5x to 91.3% at a sequencing depth of 2x. The mismatches were mainly in only one of the alleles, with <1% of the sites with both alleles imputed incorrectly. A larger number of errors were observed for heterozygous positions, mainly at a sequencing depth of 0.5x, with 27.5% of positions being miscalled with one wrong allele. The percentage of mismatches decreased up to 11.8% at a sequencing depth of 2x.

**Fig. 3.**
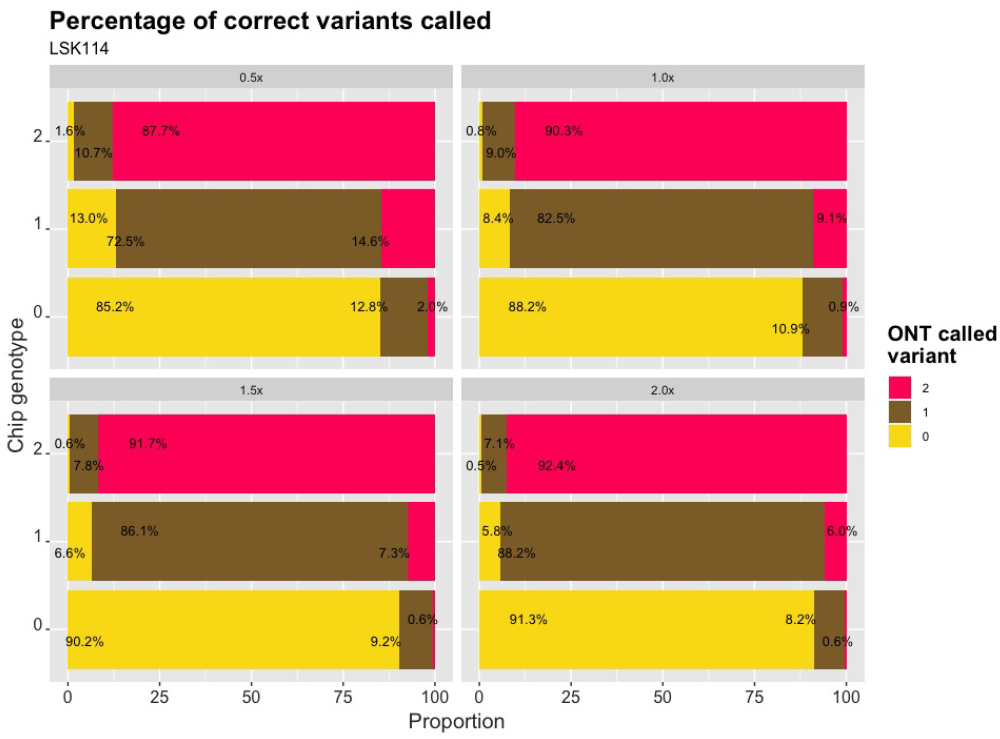
Proportion of called genotypes from LPS for each genotype code from SNP chip (vertical axes). Values are obtained from LSK114 kit at different sequencing depths (0.5x, 1.0x, 1.5x and 2.0x)

### I. Accuracy of estimated polygenic values

Global correlation between DGV calculated from SNP chips and LPS was 0.95, 0.84, and 0.95 for MY, FY and PY, respectively. However, the DGV accuracy was largely influenced by the sequencing kit. LSK114 yielded better *R*^2^ for all traits (0.92, 0.79 and 0.99 for MY, FY and PY) whereas older chemistry LSK109 showed *R*^2^ of 0.70, 0.42 and 0.58, respectively (Figures 4, 5 and 6). Regression coefficient was equal to 1 for MY using kit LSK114, and for PY using Q20 kit. Lower agreement between SNP chips and LPS was observed for FY, probably because the dispersion of this trait in the sample set was lower than for the other traits. Despite of the general strong agreement, ONT sequencing underestimated DGVs for all traits analyzed. The results from LSK109 and Q20 are comparable to the only previous study, to best of our knowledge, using ONT sequencing in a genomic selection framework: (7). However, samples in (7) were sequenced with LSK109 and at a much larger depth than in the present study, with an average yield of 22.57 Gb, which is equivalent to >7x sequence depth. DGV accuracy for samples sequenced with LSK114 was similar to those from full coverage in (7). Based on the results of our study, and in comparison to (7), the new chemistry LSK114 may provide similar accuracy as the old chemistry **but with a sequencing depth as low as 2x**.

**Fig. 4.**
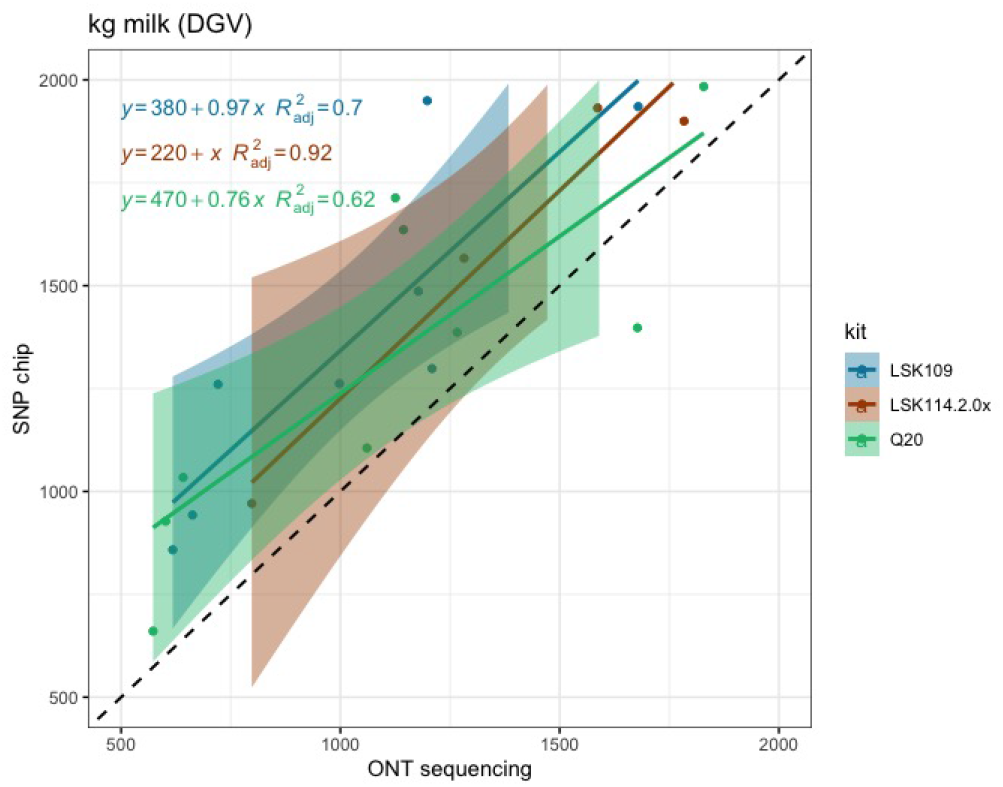
Scatter plot between Milk Yield DGVs obtained from SNP chips (y axis) and genotype-by-LPS (x axis) for the different ONT kits evaluated.

**Fig. 5.**
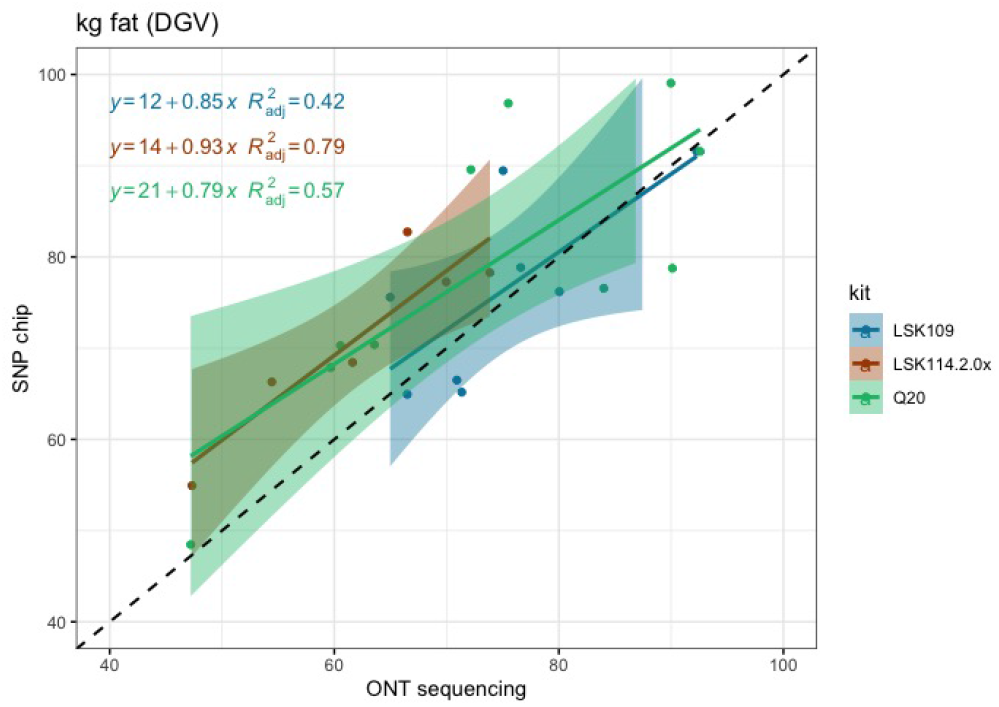
Scatter plot between Fat Yield DGVs obtained from SNP chips (y axis) and genotype-by-LPS (x axis) for the different ONT kits evaluated.

**Fig. 6.**
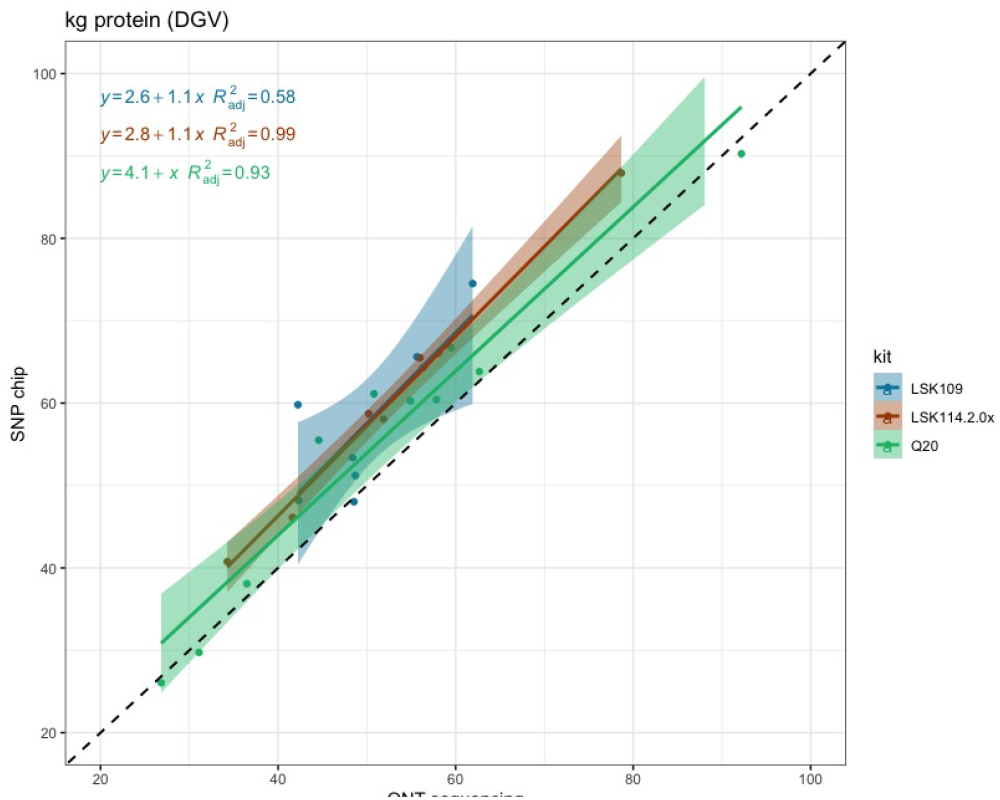
Scatter plot between DGVs obtained from SNP chips (y axis) and genotypeby-LPS (x axis) for the different ONT kits evaluated.

### I.1. Effect of sequencing depth on accuracy

Since LSK114 was the best kit performing best, we hypothesized that this might be due to a larger sequencing depth. Hence, we evaluated whether the higher DGV estimation accuracy estimated for LSK114 was due to the higher sequencing depth in Kit14 or to higher basecalling accuracies. The MY trait was used in this case. The process consisted on randomly selecting a given number of reads for each sample sequenced with the LSK114, to achieve different sequencing depths (i.e. 0.5x, 1.0x, 1.5x and 2.0x). Results are depicted in Figure 7. The *R*^2^ ranged between 0.93-0.94 for sequencing depth < 2x and 0.98 for sequencing depth of 2x. Lower sequencing depth re-sulted in more biased estimates, which may be the reason of the underestimation of the DGVs mentioned above. Larger sequencing depths (2x) alleviated this bias in the regression parameter and intercept estimation. (7) also showed more biased estimates at lower sequencing depth, and this bias was trait-dependent.

**Fig. 7.**
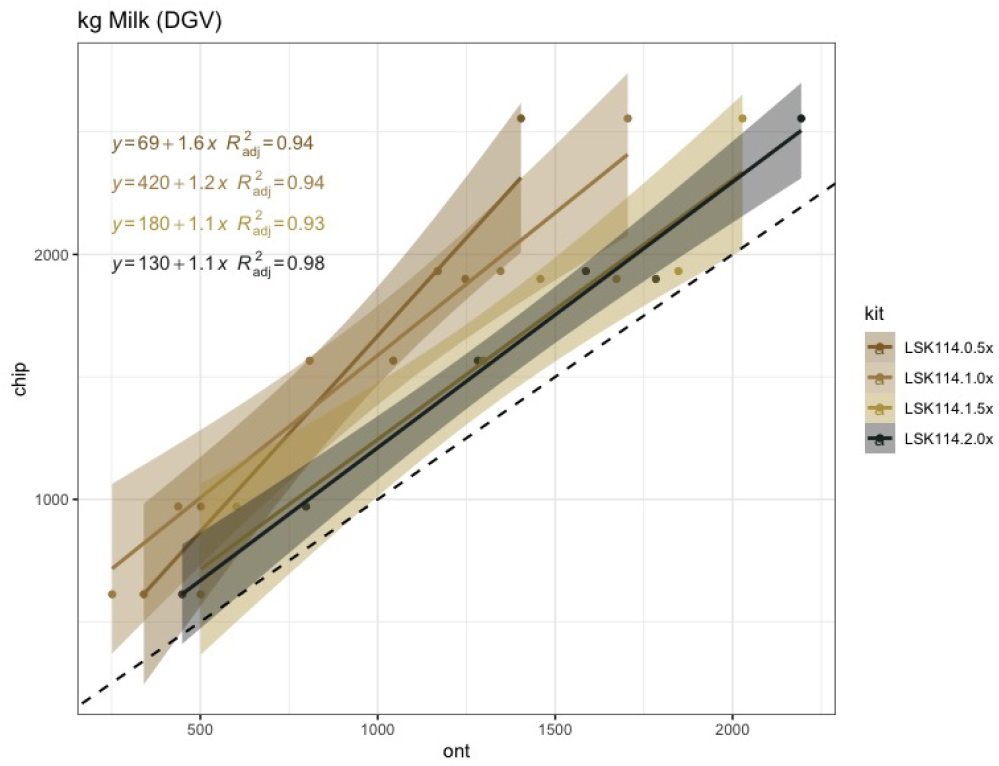
Scatter plot between Milk Yield DGVs obtained from SNP chips (y axis) and genotype-by-LPS (x axis) obtained from LSK114 kit at different sequncing depths (0.5x, 1.0x, 1.5x and 2.0x).

Table 2 shows the Spearman (rank) correlations between DGVs calculated with SNP chips and LPS. Larger correlations were calculated for LSK114 (0.94, 0.83 and 0.95), suggesting very similar ranking between SNP chips and LPS. It must be pointed out that the small samples size may negatively impact Spearman correlation. Although perfect rank agreement was not achieved in this small data set, the rank agreement obtained is encouraging to pursue new analytical methods with a large data set that may show even larger agreement for genotypes obtained by LPS.

**Table 2.**
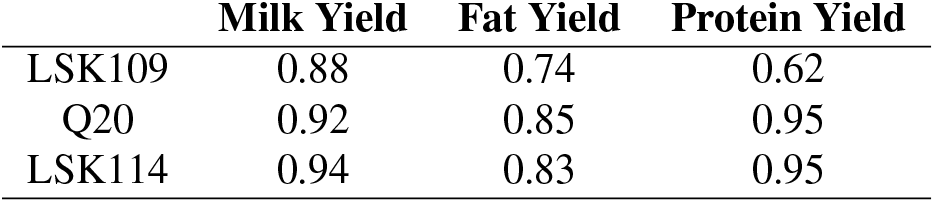
Spearman correlations between DGVs calculated from SNP chips and Nanopore genotype-by-LPS for each sequencing kit and trait evaluated

|  | **Milk Yield** | **Fat Yield** | **Protein Yield** |
| --- | --- | --- | --- |
| LSK109 | 0.88 | 0.74 | 0.62 |
| Q20 | 0.92 | 0.85 | 0.95 |
| LSK114 | 0.94 | 0.83 | 0.95 |

### J. Detection of modified bases

An average of 791 millions 5mC modifications were detected from LSK114 kit. However, after filtering for variant coverage ≥ 4x, the average amount of 5mC detected was 15.7 millions, and decreased to 2.3 and 1.6 millions for variant coverage filters ≥ 7x and ≥ 10x, respectively (Figure 8). In terms of sequencing yield, 5-6 Gb would produce more than 15 million 5mC methylation states at a coverage ≥ 4x, and at least 1.5 million 5mC sites at coverage ≥ 10x. We evaluated the differences for coverage filters of 4x, 7x and 10x. A large agreement in the methylation percentage was observed in genomic bins of 500 bp: a correlation of 0.985 was achieved between filters ≥ 4x and ≥ 10x, and 0.986 between ≥ 7x and ≥ 10x. Figure 12 depicts the genome positions that were methylated for each sample sequenced with the LSK114 kit after filtering for coverage ≥ 4x. Methylation was detected along the whole genome. Samples with a larger genome coverage and sequencing yield (samples 25, 26 and 30) also showed a larger density of methylated position across the genome at a coverage ≥ 4x. Those samples with lower genome coverage still showed a genome-wide methylation randomly distributed along the genome, although with a lower density of the methylated sites. Filtering for coverage ≥ 10x led to a much sparse methylation marks, being critical for those samples with lower sequencing depth.

**Fig. 8.**
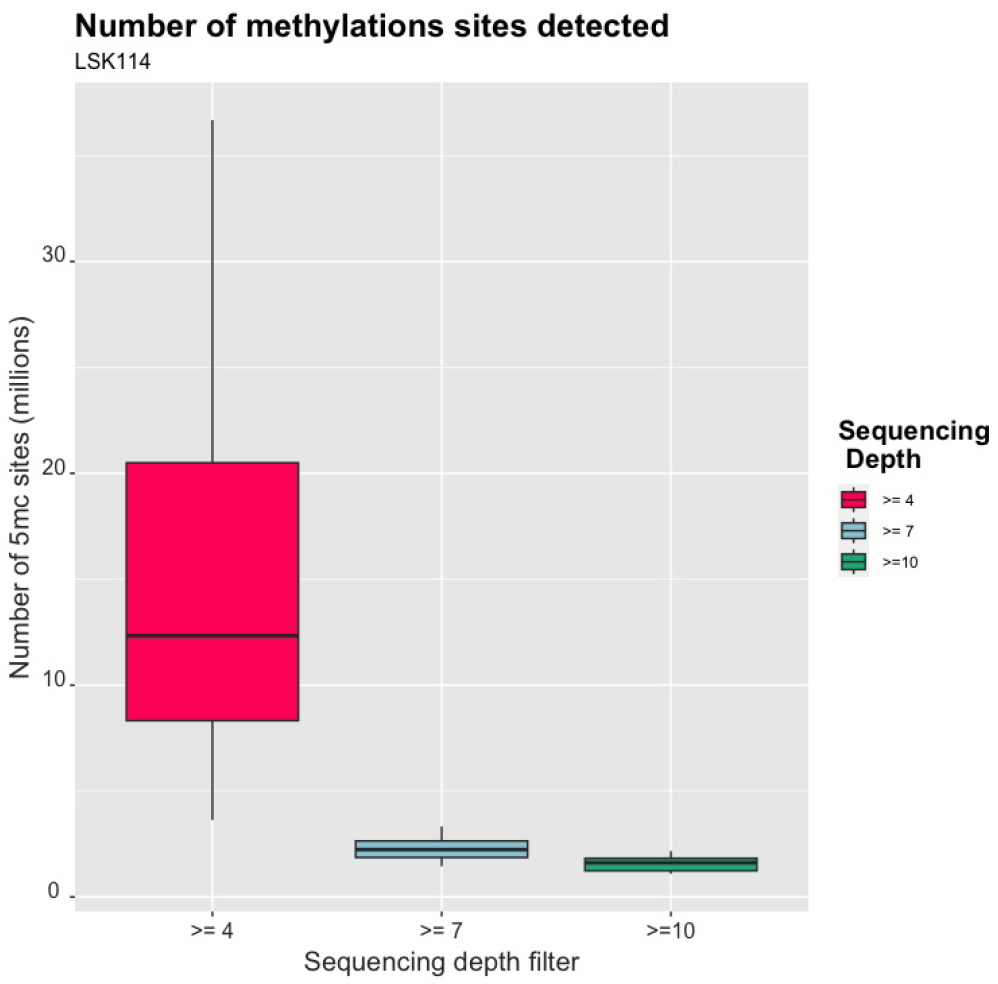
Boxplots for the number of methylated sites obtained from LPS with LSK114 kit after filtering by sequencing depth ≥ 4, ≥ 7 or ≥ 10. Global sequencing depth from LPS was 2x.

These methylation marks were located mainly in distal intergenic regions (Figure 10), emphasizing the evidence that the genome is pervasively transcribed, and that the majority of its bases are in primary transcripts, including non-protein-coding transcripts (16). Around 5-6 % of methylated positions were found in promoter regions, and there were little variability in this percentage among samples. Larger variability was found in the percentage of methylated sites found in exons and distal intergenic regions. Filtering for coverage ≥ 10x led to similar proportions at promoter regions, but a much larger proportion of methylated sites in distal intergenic regions (Figure 11). This implies that too stringent coverage filters applied to LPS may under-represent methylation in promoter regions, since many methylated sites are filtered out. After filtering for coverage ≥ 4x, the methylation pattern was as expected with a larger density of methylation marks at TSS, and a sudden drop upstream (Figure 9). Figure 9 shows the methylation status near the TSS of known genes. Some genes showed large proportion of methylation marks at or near-by the TSS, which is often maintained upstream during few hundreds bases. Interestingly, other genes showed no methylation at the TSS or nearby, probably because they are constitutive or necessary genes. This deserves further study. Therefore, a minimum coverage of 4x may be enough for whole genome methylation, considering that it showed very similar results and high accordance with a coverage filter ≥ 10x, and kept 10-folds more sites.

**Fig. 9.**
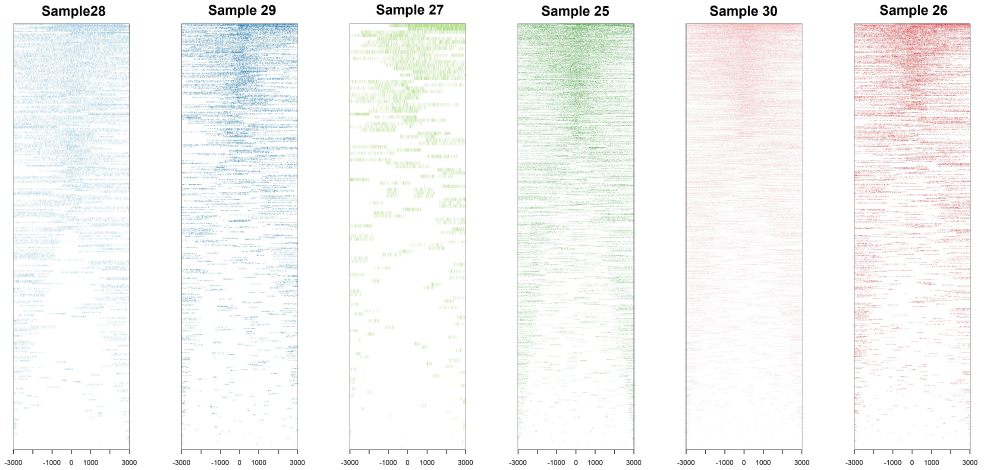
Density of methylated position found for each sample according to their distance to the Transcription Start Site (TSS). Samples were sequenced with LSK114 kit. The plot shows the position left after filtering by sequencing depth ≥ 4. Global sequencing depth from LPS was 2x.

**Fig. 10.**
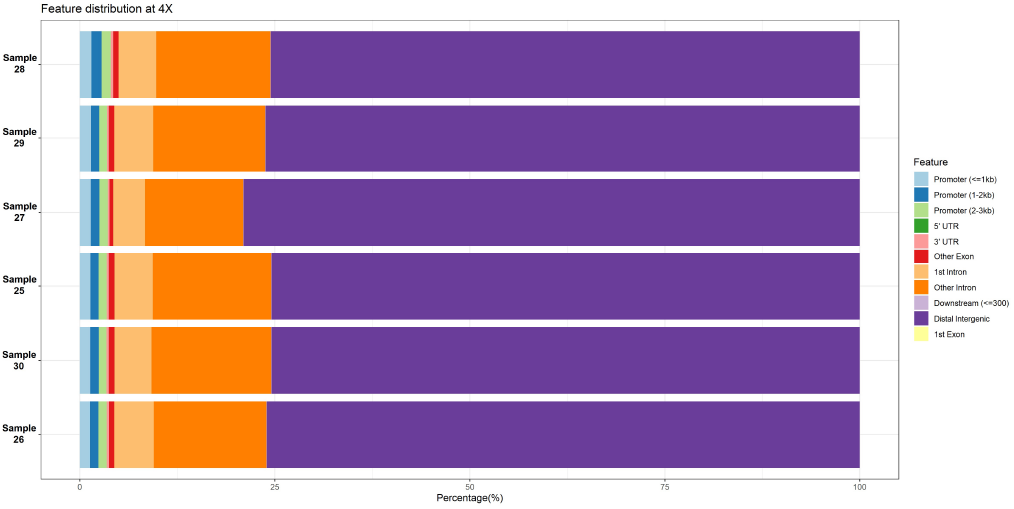
Percentage of methylated genomic regions found for each sample sequenced with the LSK114 kit. Positions showed had a coverage ≥ 4x.

**Fig. 11.**
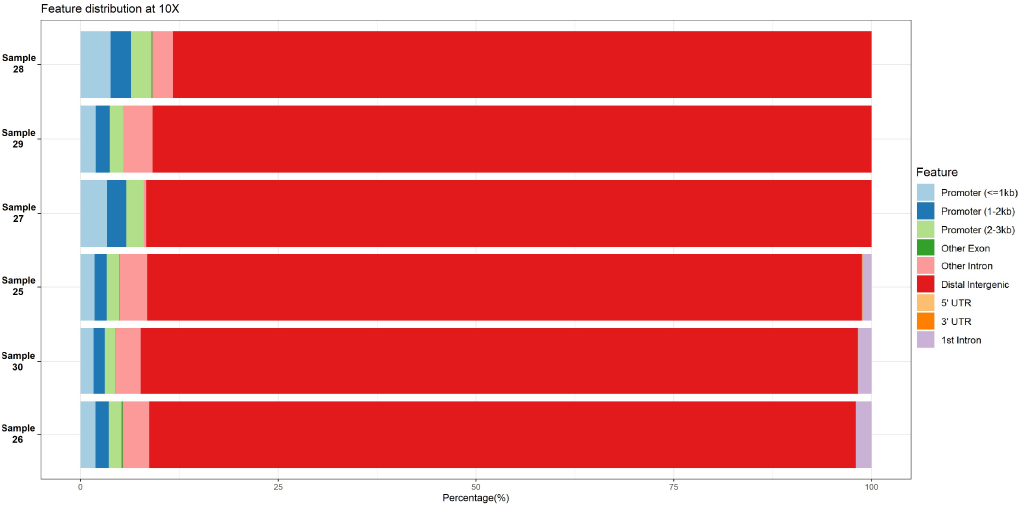
Percentage of methylated genomic regions found for each sample sequenced with the LSK114 kit. Positions showed had a coverage ≥ 10x.

**Fig. 12.**
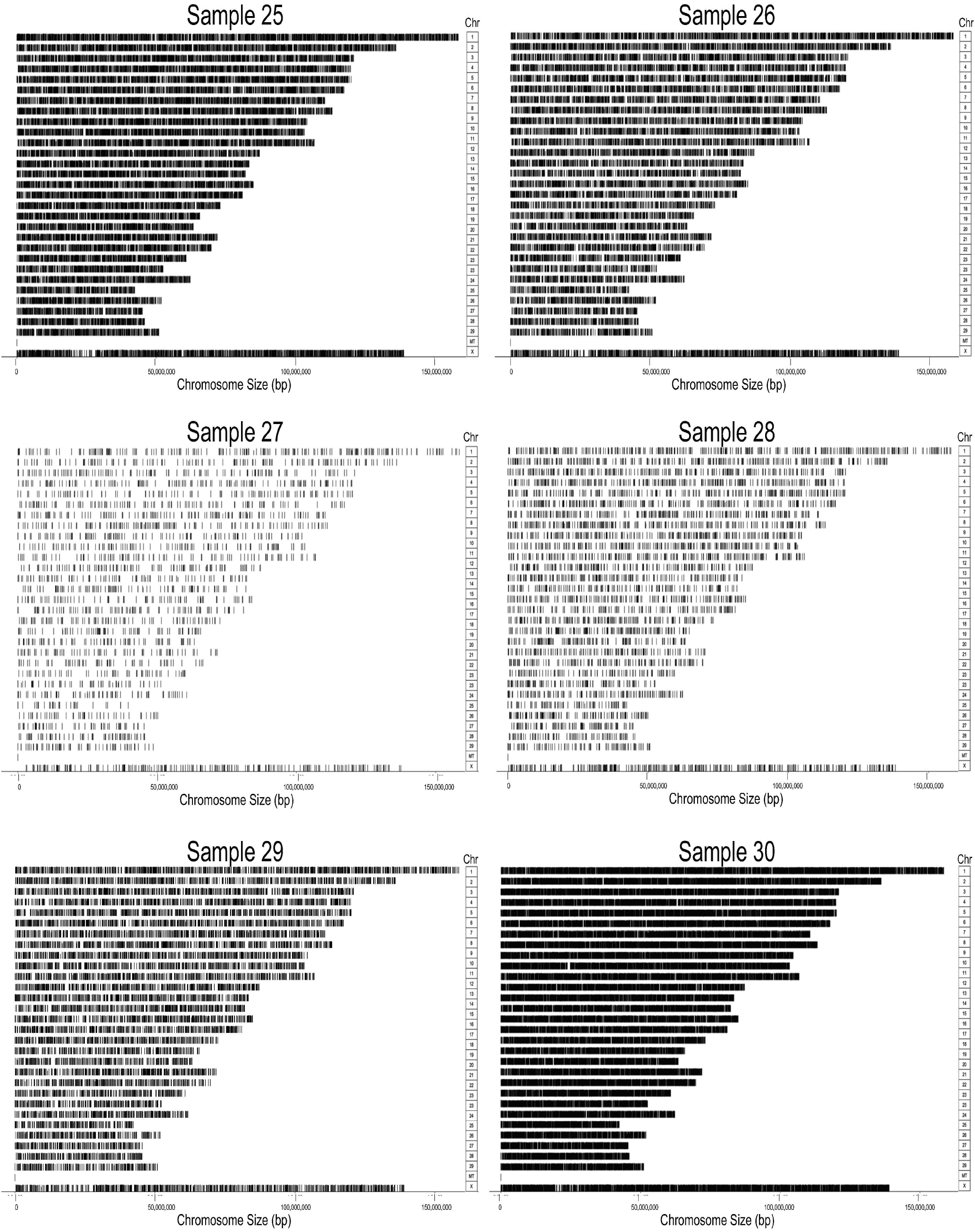
Chromosome-wide methylation sites for each sample sequenced with the LSK114 kit. Positions showed had a coverage ≥ 4x.

## Conclusions

This is the first study evaluating the DGV accuracy obtained from different ONT chemistries in a genotype-by-LPS framework, and simultaneously extracting methylation marks. Older ONT chemistry still pose some bias when used at low sequencing depth. However, the latest LSK114 chemistry provided a high basecalling accuracy that was suitable for breeding value prediction in a genomic selection framework, in spite of a lower imputation accuracy achieved on heterozygote genotypes. This limitation may be alleviated by using DA to estimate DGV or polygenic risk scores at a low sequencing depth of 2x. Lower sequencing depths may still provide high ranking agreement yet with larger bias. The study showed the possibility to obtain methylation status throughout the genome with a high reliability even at a low sequencing depth, which comes at no extra cost with genotype-by-LPS. This epigenetic information can be used in epigenomic wide association studies to infer association between methylation and phenotypic expression of traits of interest. It can also be included in mixed models to account for epigenetic variance or determine the effect of environmental forces on the methylation status. Despite its affordability, it is not expected that genotype-by-LPS can compete with SNP genotyping arrays at its current cost. In the future, an increased base calling accuracy may lead genotype-by-LPS to achieve similar DGV accuracy as SNP genotypes, even at lower sequencing depths. Proper multiplexing strategies may contribute to decrease the cost. Genotype-by-LPS is also appealing in populations where SNP chips are not available or a high density of markers is required (e.g. populations with short linkage disequilibriun range). New research and field application opportunities arise with the proposed genotype-by-LPS in livestock breeding programs and also at evaluating management practices that may impact on the epigenetic status of the animals. Our results showed that genotype-by-LPS is attractive for research including genomic and epigenomic variants, despite of few limitations such as a lack of full agreement with SNP chip genotypes, and low coverage of methylation marks.

## ACKNOWLEDGEMENTS

The Spanish Holstein Association (CONAFE) is acknowledge for providing SNP chip genotypes and allele substitution effects to calculate direct genomic values.

